# Comparative genomics of *Bordetella pertussis* isolates from New Zealand, a country with an uncommonly high incidence of whooping cough

**DOI:** 10.1101/2021.10.11.463938

**Authors:** Natalie Ring, Heather Davies, Julie Morgan, Maithreyi Sundaresan, Audrey Tiong, Andrew Preston, Stefan Bagby

## Abstract

2.

Whooping cough, the respiratory disease caused by *Bordetella pertussis*, has undergone a wide-spread resurgence over the last several decades. Previously, we developed a pipeline to assemble the repetitive *B. pertussis* genome into closed sequences using hybrid nanopore and Illumina sequencing. Here, this sequencing pipeline was used to conduct a more high-throughput, longitudinal screen of 66 strains isolated between 1982 and 2018 in New Zealand. New Zealand has a higher incidence of whooping cough than many other countries; usually at least twice as many cases per 100,000 people as the USA and UK and often even higher, despite similar rates of vaccine uptake. To the best of our knowledge, these strains are the first New Zealand *B. pertussis* isolates to be sequenced. The analyses here show that, on the whole, genomic trends in New Zealand *B. pertussis* isolates, such as changing allelic profile in vaccine-related genes and increasing pertactin deficiency, have paralleled those seen elsewhere in the world. At the same time, phylogenetic comparisons of the New Zealand isolates with global isolates suggest that a number of strains are circulating in New Zealand which cluster separately from other global strains, but which are closely related to each other. The results of this study add to a growing body of knowledge regarding recent changes to the *B. pertussis* genome, and are the first genetic investigation into *B. pertussis* isolates from New Zealand.

**Impact statement:** Since the 1990s, whooping cough has been resurgent in many countries around the world, despite the wide availability of vaccines. New Zealand has often had a higher incidence of whooping cough than other countries such as the USA, UK and Australia, both during outbreak periods and in the intervening years. One potential reason for the resurgence of whooping cough is genetic changes to the causative bacterium, *Bordetella pertussis*, with several recently identified, ongoing global genetic trends. No *B. pertussis* isolates from New Zealand have previously been sequenced, however. Here, we used hybrid sequencing to investigate the genomes of 66 New Zealand *B. pertussis* isolates, collected between 1982 and 2018. This revealed that genetic trends in New Zealand *B. pertussis* match those observed elsewhere, but over the years a number of highly similar or identical strains appear to have circulated (or are currently circulating) in the country, a phenomenon not commonly noted elsewhere. This first study of *B. pertussis* isolates from New Zealand contributes to the global understanding of *B. pertussis* genomics, as well as providing the groundwork for any future whole genome sequencing of New Zealand *B. pertussis* isolates.

**Data summary:**

1. Nanopore and Illumina fastq sequence files for all strains have been deposited in NCBI’s Sequence Read Archive, BioProject PRJNA556977. A full list of accession numbers for all sequence read files is provided in **Supplementary Table S1**.
2. Genome sequences for 63 strains have been deposited in NCBI’s GenBank, BioProject PRJNA556977, accession numbers in **Supplementary Table S1**.
3. Genome assemblies for 3 strains assembled using only nanopore data (NZ1, NZ5 and NZ29), which had a high number of pseudogenes, were not deposited in GenBank, but are available from Figshare: https://doi.org/10.6084/m9.figshare.12640463
4. Source code and full commands used are available from Github: https://github.com/nataliering/Comparative-genomics-of-Bordetella-pertussis-isolates-from-New-Zealand

The authors confirm all supporting data, code and protocols have been provided within the article or through supplementary data files.

## 5. Introduction

Despite the availability of vaccination, whooping cough, the respiratory disease caused by *Bordetella pertussis*, has been resurgent in many countries for the past several decades. Proposed causes of this resurgence include changes to the bacterium at the genomic level, potentially in response to the introduction of an acellular whooping cough vaccine (ACV) in the late 1990s (1). The ACV contains one to five *B. pertussis* antigens, including Pertussis toxin (Ptx), pertactin (Prn), filamentous haemagluttinin (FHA), and the fimbrial proteins Fim2 and Fim3. Whooping cough vaccines have been available in New Zealand since 1945, and part of the immunisation schedule since 1960. New Zealand switched from the first generation whole cell vaccines (WCV) to a three-antigen (Ptx-FHA-Prn) ACV in 2000.

Extensive screens of strains circulating during and between epidemics in the UK, Australia and the USA have been conducted in recent years (2–7), each contributing to our understanding of how *B. pertussis* is evolving under selection pressure from vaccines. Key observations from these studies include a shifting allelic profile in many of the genes encoding ACV antigens, such as the Pertussis toxin promoter, ptxP. A landmark 2014 study by Bart et al. (7) showed that, globally, the alleles for the ACV genes have been changing since the introduction of the WCV, and ostensibly faster since the switch to the ACV in many countries. A subsequent study of strains from a 2012 whooping cough outbreak in the UK confirmed that the ACV genes do indeed appear to be mutating at a faster rate than the genes which code for other cell surface proteins (2).

Another, more recently observed, trend in the *B. pertussis* genome is the increasing prevalence of strains which are deficient for expression of one or more of the ACV antigens. Prior to the introduction of the ACV, only a handful of strains had been identified which possessed mutations resulting in their inability to express functional Prn (8–10). Bart et al.’s 2014 study, for example, included 323 strains which were all isolated prior to 2010, and none of these were found to be pertactin deficient. Since the mid-2000s, however, an increasing number of pertactin deficient strains have been observed in many countries in the world. In Australia, the percentage of isolates found to be pertactin deficient increased from 5 to 78% between 2008 and 2012 (3). Likewise, 640 of 753 strains isolated in the US between 2011 and 2013 were pertactin deficient (11). A longitudinal study of European *B. pertussis* isolates found a correlation between the length of time a country had been using the ACV, and the percentage of pertactin deficient isolates from that country (12). A number of different mutations resulting in pertactin deficiency have been identified as including deletions, SNPs and the insertion of the abundant *B. pertussis* insertion sequence, IS *481* into various loci within the pertactin gene, *prn*, although no single mechanism for deficiency appears to be predominant (12–14). This suggests a selection for loss of expression of pertactin, rather than expansion of a single pertactin-deficient lineage.

A final genomic phenomenon which has become apparent over the last decade is the existence of a large number of different genome arrangements in circulating *B. pertussis* isolates. There is some evidence, such as the work of Dienstbier et al. (15), that different genomic arrangements have different transcriptomic profiles. Isolates with different genomic arrangements may therefore also vary phenotypically. Work in this area is ongoing, and the exact contribution, if any, of genomic arrangement to phenotypes in *B. pertussis* is still poorly understood. However, in recent years, long-read sequencing technologies, such as Pacific Biosciences and Oxford Nanopore Technologies, have enabled large-scale studies into *B. pertussis* genome arrangement, using short- and long-read technologies in tandem to assemble closed genome sequences. These have produced a catalogue of existing arrangements, and their prevalence, to which the closed genome sequences of future isolates can be compared (5, 16, 17).

Since the whooping cough vaccine was introduced globally, New Zealand has noted a high number of infections and hospitalisations due to whooping cough compared with other developed countries; some sources indicate disease rates 5 to 10 times higher in New Zealand than the UK or USA. For instance, during the 1980s, hospitalisation for whooping cough was required for 0.37 per 100,000 people in the USA, compared with 3.75 per 100,000 in New Zealand (18). In addition, epidemic periods have often been more severe in New Zealand than other countries. During concurrent epidemics in the UK and New Zealand in 2012, 122.3 cases were seen per 100,000 people in New Zealand, compared to 20 per 100,000 in the UK (2, 19). Whilst, historically, rates of vaccine uptake in New Zealand were relatively low, a marked increase in immunisation coverage across all age groups has been seen over the last two decades (20, 21). At 93%, immunisation coverage in New Zealand at all age points up to 5 years is now comparable to the mean 94% coverage in the United Kingdom (22). Nonetheless, rates of whooping cough have remained higher in New Zealand than elsewhere, with some outbreaks occurring which are unique to New Zealand. The latest was announced in December 2017 (23). By May 2019, notifications of whooping cough cases had returned to the “expected” level, after a total of 4,697 cases, or around 96 per 100,000 people across the 19-month outbreak period (24).

As of 2018, no *B. pertussis* sequencing reads or genomes in the NCBI database were tagged as being from New Zealand. However, *B. pertussis* isolates have been collected and stored at the Institute of Environmental Science and Research (ESR)’s Invasive Pathogen Laboratory in New Zealand since the 1980s (25). Here, we sequenced the genomes of 66 *B. pertussis* isolates collected in New Zealand between 1982 and 2018, using a variant of the nanopore-based hybrid sequencing workflow we developed previously (26). We then used comparative genomics to place these first New Zealand *B. pertussis* genome sequences in a global context, and to screen for any potential genetic factors responsible for New Zealand’s ongoing high incidence of whooping cough.

## 6. Methods

All data processing and analysis was carried out using the MRC’s Cloud Infrastructure for Microbial Bioinformatics (CLIMB) (27).

### Strain isolation

66 strains, collected between 1982 and late 2018, were stored at −80°C at the New Zealand Institute of Environmental Science and Research (ESR)’s Kenepuru Science Centre, Porirua, New Zealand. Strains were grown and heat-killed, then shipped on ice to the University of Bath, United Kingdom. On arrival, the heat-killed cells were stored at −20°C. Serotyping data had previously been determined by the ESR as the strains were isolated, where possible (46/66 strains). Full details, including accession numbers, are included in **Table 1** and **Supplementary Table S1**.

**Table 1.** Details of the New Zealand isolates sequenced here. Further details can be found in **Supplementary Table S1**.

| Isolate | Year | Serotype | *ptxP* | *ptxA* | *fim2* | *fim3* | *prn* | *prn* mutation | Arrangement type |
| --- | --- | --- | --- | --- | --- | --- | --- | --- | --- |
| NZ1 | 1982 | Unknown | 1 | 1 | 1 | 1 | 1 | NA | NZ006 |
| NZ2 | 1983 | Unknown | 1 | 1 | 1 | 1 | 1 | NA | NZ006 |
| NZ3 | 1984 | Unknown | 1 | 1 | 1 | 1 | 1 | NA | NZ006 |
| NZ4 | 1985 | Unknown | 1 | 1 | 1 | 1 | 1 | NA | Singleton |
| NZ5 | 1986 | Unknown | 1 | 1 | 1 | 1 | 1 | NA | Singleton |
| NZ6 | 1987 | Unknown | 3 | 1 | 1 | 1 | 2 | NA | CDC010 |
| NZ7 | 1988 | Unknown | 1 | 1 | 1 | 1 | 9 | NA | Singleton |
| NZ8 | 1989 | 1,3 | 3 | 1 | 1 | 1 | 2 | NA | CDC010 |
| NZ9 | 1990 | 1,3 | 3 | 1 | 1 | 1 | 2 | NA | CDC002 |
| NZ10 | 1991 | 1,2 | 1 | 1 | 1 | 1 | 1 | NA | Singleton |
| NZ11 | 1993 | 1,3 | 3 | 1 | 1 | 1 | 2 | NA | CDC010 |
| NZ12 | 1994 | 1,3 | 1 | 1 | 1 | 1 | 3 | NA | Singleton |
| NZ13 | 1997 | 1,3 | 1 | 1 | 1 | 1 | 3 | NA | NZ002 |
| NZ14 | 1999 | Unknown | 3 | 1 | 1 | 1 | 2 | NA | CDC002 |
| NZ15 | 2000 | Unknown | 3 | 1 | 1 | 2 | 2 | NA | CDC013 |
| NZ16 | 2001 | Unknown | 3 | 1 | 1 | 2 | 2 | NA | CDC082 |
| NZ17 | 2002 | 1,3 | 3 | 1 | 1 | 1 | 2 | NA | CDC010 |
| NZ18 | 2002 | 1,3 | 3 | 1 | 1 | 2 | 2 | NA | CDC013 |
| NZ19 | 2003 | 1,3 | 3 | 1 | 1 | 2 | 2 | NA | CDC013 |
| NZ20 | 2007 | 1,3 | 3 | 1 | 1 | 2 | 2 | NA | NZ007 |
| NZ21 | 2007 | 1,3 | 3 | 1 | 1 | 1 | 2 | NA | CDC010 |
| NZ22 | 2013 | 1,2 | 3 | 1 | 1 | 1 | 2 | prn2::IS481-1613fwd | CDC010 |
| NZ23 | 2013 | 1,2,3 | 3 | 1 | 1 | 1 | 2 | prn2::IS481-1613rev | CDC010 |
| NZ24 | 2014 | Unknown | 3 | 1 | 1 | 1 | 2 | prn2::IS481-1613rev | CDC010 |
| NZ25 | 2014 | 1, 2 | 3 | 1 | 1 | 1 | 2 | prn2::IS481-1613rev | CDC010 |
| NZ26 | 2015 | 1, 2 | 3 | 1 | 1 | 1 | 2 | NA | CDC002 |
| NZ27 | 2015 | 1, 3 | 3 | 1 | 1 | 1 | 2 | prn2::IS481-1613fwd | CDC010 |
| NZ28 | 2016 | 1, 2, 3 | 3 | 1 | 1 | 2 | 2 | NA | Singleton |
| NZ29 | 2004 | 1,3 | 3 | 1 | 1 | 1 | 1 | NA | CDC010 |
| NZ30 | 2004 | 1,3 | 3 | 1 | 1 | 1 | 2 | NA | CDC010 |
| NZ31 | 2004 | 1,3 | 3 | 1 | 1 | 1 | 2 | NA | CDC010 |
| NZ32 | 2005 | 1,2,3 | 1 | 1 | 1 | 1 | 2 | NA | Singleton |
| NZ33 | 2005 | 1,3 | 3 | 1 | 1 | 1 | 2 | NA | CDC010 |
| NZ34 | 2005 | 1,3 | 3 | 1 | 1 | 1 | 2 | NA | CDC010 |
| NZ35 | 2005 | 1,2 | 1 | 1 | 1 | 1 | 3 | NA | NZ002 |
| NZ36 | 2006 | 1,2 | 1 | 1 | 1 | 1 | 3 | NA | NZ002 |
| NZ37 | 2006 | 1,2,3 | 1 | 1 | 1 | 1 | 3 | NA | NZ002 |
| NZ38 | 2008 | 1,3 | 3 | 1 | 1 | 1 | 2 | NA | CDC010 |
| NZ39 | 2008 | 1,3 | 3 | 1 | 1 | 2 | 2 | NA | NZ007 |
| NZ40 | 2009 | 1,3 | 3 | 1 | 1 | 2 | 2 | NA | CDC082 |
| NZ41 | 2009 | 1,2 | 1 | 1 | 1 | 4 | 1 | NA | Singleton |
| NZ42 | 2009 | 1,3 | 3 | 1 | 1 | 1 | 2 | NA | CDC010 |
| NZ43 | 2010 | 1,3 | 3 | 1 | 1 | 1 | 2 | NA | CDC010 |
| NZ44 | 2010 | 1,3 | 3 | 1 | 1 | 1 | 2 | NA | NZ008 |
| NZ45 | 2011 | 1,3 | 3 | 1 | 1 | 2 | 2 | NA | CDC082 |
| NZ46 | 2011 | 1,2 | 3 | 1 | 1 | 1 | 2 | NA | CDC002 |
| NZ47 | 2012 | 1,2 | 3 | 1 | 1 | 1 | 2 | prn2::IS481-1613fwd | CDC010 |
| NZ48 | 2012 | 1,3 | 3 | 1 | 1 | 2 | 2 | prn2::IS481-1613rev | CDC046 |
| NZ49 | 2012 | 1,3 | 3 | 1 | 1 | 1 | 2 | prn2::IS481-240rev | CDC002 |
| NZ50 | 2012 | 1,3 | 3 | 1 | 1 | 1 | 2 | prn2::IS481-1613fwd | CDC010 |
| NZ51 | 2012 | 1,3 | 3 | 1 | 1 | 1 | 2 | prn2::IS481-1613fwd | CDC010 |
| NZ52 | 2012 | 1,3 | 3 | 1 | 1 | 1 | 2 | prn2::IS481-1613rev | CDC010 |
| NZ53 | 2012 | 1,3 | 3 | 1 | 1 | 1 | 2 | prn2::IS481-1613fwd | CDC010 |
| NZ54 | 2012 | 1,2 | 3 | 1 | 1 | 1 | 2 | prn2::IS481-1613rev | CDC010 |
| NZ55 | 2012 | 1,3 | 3 | 1 | 1 | 1 | 2 | prn2::IS481-1613rev | CDC010 |
| NZ56 | 2012 | 1,3 | 3 | 1 | 1 | 1 | 2 | prn2::IS481-1613rev | CDC010 |
| NZ57 | 2017 | Unknown | 3 | 1 | 1 | 1 | 2 | prn2::IS481-1613fwd | CDC237 |
| NZ58 | 2017 | Unknown | 3 | 1 | 1 | 1 | 2 | prn2::IS481-1613fwd | CDC237 |
| NZ59 | 2017 | Unknown | 3 | 1 | 1 | 1 | 2 | prn2::IS481-1613fwd | CDC010 |
| NZ60 | 2017 | Unknown | 3 | 1 | 1 | 1 | 2 | prn2::IS481-1613fwd | NZ008 |
| NZ61 | 2017 | Unknown | 3 | 1 | 1 | 1 | 2 | prn2::IS481-1613fwd | CDC010 |
| NZ62 | 2017 | 1, 2 | 3 | 1 | 1 | 1 | 2 | prn2::IS481-1613fwd | CDC010 |
| NZ63 | 2017 | Unknown | 3 | 1 | 1 | 1 | 2 | prn2::Stop-C223T | Singleton |
| NZ64 | 2018 | Unknown | 3 | 1 | 1 | 1 | 2 | prn2::IS481-1613fwd | CDC237 |
| NZ65 | 2018 | Unknown | 3 | 1 | 1 | 1 | 2 | prn2::IS481-1613fwd | CDC237 |
| NZ66 | 2018 | Unknown | 3 | 1 | 1 | 1 | 2 | NA | CDC010 |

### DNA extraction and Illumina sequencing

Heat-killed cells were resuspended in 1 ml PBS and OD_600_ was measured. Volumes of suspension equating 1ml at an OD_600_ of 2.0 (~4×10^9^ *B. pertussis cells*) were pelleted in a microcentrifuge for 2 min at 12,000 xg. gDNA was extracted from each pellet using the QIAamp DNA mini kit (Qiagen) according to the manufacturer’s instructions, including a single-step elution into 200 μl of elution buffer (buffer AE).

gDNA from 34 isolates was sent for Illumina MiSeq sequencing at the Milner Centre, University of Bath. gDNA from the remaining 32 isolates was sent to Novogene for Illumina NovaSeq sequencing. gDNA from 3 isolates was lost during shipping, hence 63 isolates were sequenced with Illumina sequencers.

### Nanopore library preparation and sequencing

Sequencing libraries were prepared for all samples using ONT’s Rapid Barcoding Kit (SQK-RBK004), according to the manufacturer’s instructions, including the optional 1x SPRI concentration and clean-up step (using Promega ProNex size selection beads) after library pooling and before addition of RAP adaptors. Between 10 and 12 barcodes were used per MinION flow cell (see **Supplementary Table S2** for full details).

Each pooled sequencing library was loaded onto an R9.4 MinION flow cell and sequenced for 48 h using a MinION Mk1b device with MinKNOW sequencing software.

### Basecalling, demultiplexing and adaptor trimming

Deepbinner (v0.2.0, 28) was used to demultiplex the raw fast5 files, using the “realtime” setting with “rapid” option. This placed all fast5 files into separate bins, one for each barcode. ONT’s Guppy fast Flip-flop basecaller (v3.1.5+781ed57) was then run on each of these barcode bins, with its own rapid barcode demultiplexing option enabled. This placed each of the fastq files basecalled from the fast5s in the Deepbinner barcode bins into further barcode bins, resulting in twelve Deepbinner barcode bin fast5 directories (plus one “unclassified” bin, containing reads with unclassifiable barcodes), each containing up to twelve further barcode bin fastq directories (plus “unclassified”). Theoretically, most of the reads from any specific Deepbinner bin should have been placed in a bin corresponding to the same barcode by Guppy, but in some cases the two tools may disagree over the barcode identity; for example, most of the reads in the “barcode01” Deepbinner bin should have also been placed in the “barcode01” bin by Guppy after basecalling, but some may have been placed in other bins. Only reads identified as having the same barcode by both tools were retained for further processing; for example, when basecalling the fast5s from Deepbinner’s “barcode01” bin, only fastqs placed in Guppy’s “barcdoe01” bin were kept. Finally, Porechop (v0.2.4, 29) was used to trim the barcodes and other adaptor sequences from the demultiplexed reads.

### Hybrid genome assembly

Illumina MiSeq data was provided pre-trimmed by the Milner Genomics Centre. Illumina NovaSeq data provided by Novogene was trimmed using Trimmomatic (v0.36, 30) with the options PE, HEADCROP:10, SLIDINGWINDOW:4:25 and MINLEN:100. Genome assembly was attempted for all strains using the hybrid pipeline we developed previously (26). Nanopore reads were pre-corrected using Canu (v1.8, 31), followed by hybrid assembly with nanopore and Illumina reads using Unicycler (v0.4.7, 32). However, some of the available Illumina data was of lower quality (reads <70 bp, non-uniform length) or low coverage (<40x). Unicycler uses an Illumina-centric hybrid assembly method, using SPAdes (33) to first assemble the Illumina data into contigs, then the nanopore reads to attempt to bridge the contigs. The low quality of some of the Illumina data therefore prevented Unicycler from assembling closed genomes for these strains. An updated variant of the best long-read-centric hybrid assembly strategy identified in Ring et al. 2018 (26) was used to assemble genomes for the strains with low quality Illumina data (full details of which strategy was used for each strain are shown in **Supplementary Table S1**).

For the isolates which required long-read-centric assembly, Canu-corrected nanopore reads were assembled with Flye (v2.7b-b1526, 34) and polished four times with Racon (v1.4.11, 35), then polished with Medaka (v0.11.3, https://github.com/nanoporetech/medaka), both using the nanopore reads alone. Finally, the assembly was polished three times using Pilon (v1.22, 36) with the short Illumina reads.

Illumina reads were not available for three strains (NZ1, NZ5 and NZ29). Genome assemblies were produced for these strains using nanopore reads alone, using the Flye-Racon-Medaka strategy above, without the Pilon polishing steps. These strains were not included in SNP analysis, phylogenies or allele typing, but they were included in analysis of genome arrangement and numbers of IS elements.

All sequences were rearranged to start with *gidA*, and reverse complemented where necessary, as described previously (26).

### Comparing genome arrangement

The closed genome sequences described above were aligned against each other using progressiveMauve (v20150226 build 10, 37). The results were manually inspected and grouped into types (within each type, no arrangement differences were visible). A representative of each New Zealand arrangement type was aligned against 29 global strains (**Supplementary Table S3**), representing each of the CDC arrangement types defined in Weigand, Peng (17)’s landmark *B. pertussis* genomic arrangement study, in order to place the New Zealand arrangements in a wider global context.

### Allelic profile typing

Allele type was assigned to the genes coding for the ACV proteins (*ptxA-E, prn, fim2, fim3, fhaB*) and the promoter for pertussis toxin (*ptxP*) using a custom-made MLST scheme with Seemann’s MLST tool (v2.16.2, 38), using the hybrid assembled genomes. The commands and custom scheme are available from https://github.com/nataliering/Comparative-genomics-of-Bordetella-pertussis-isolates-from-New-Zealand and https://github.com/nataliering/Comparative-genomics-of-Bordetella-pertussis-isolates-from-New-Zealand, respectively. Instructions for using a custom scheme are available from https://github.com/tseemann/mlst. To make the ACV-gene MLST scheme, all known alleles were downloaded for each gene from PubMLST (39) and processed using tfa_prepper (https://github.com/nataliering/Comparative-genomics-of-Bordetella-pertussis-isolates-from-New-Zealand). The exceptions were *prn* and *fim3*, for which the alleles defined in Bart, Harris (7) (supplemental text sd6) were used for consistency, as the nomenclature for both on PubMLST appears to be different.

### Prediction of pertactin, pertussis toxin and filamentous haemagluttinin deficiency

The final closed genome sequence for each strain was annotated using Prokka (v1.14.6, 40), with the Tohama I reference proteins for NC_002929.2 from GenBank as a guide. The resulting annotations for *prn*, *ptxA-E* and *fhaB* were screened for presence/absence, and for insertion of IS *481*, IS *1002*, or IS *1663*. Additionally, the assembled *prn* sequences were screened for the presence of other mutations previously identified in pertactin-deficient strains (41).

### Phylogenies

To place the New Zealand strains in a global context, 73 global strains representing different continents, time periods and allelic profiles were included in the analysis. A further 125 global strains were included in more detailed trees for each of New Zealand’s four whooping cough outbreaks since 2004. Illumina reads were downloaded from NCBI’s SRA using fasterq-dump (v2.10.5) and trimmed using Trimmomatic (v0.36) using the options HEADCROP:10, SLIDINGWINDOW:4:20 and MINLEN:30 (see **Supplementary Table S4** for full details including accession numbers).

Several different trees were constructed: one containing only the New Zealand strains, one containing the New Zealand strains along with the 73 global strains, and one for each of the four New Zealand whooping cough outbreaks since 2004. Snippy (v4.6.0, 40) and SNP-sites (v2.1.3, 42) were used to perform variant calling and define the core genome, using the paired-end Illumina fastq files for all test strains, with the Tohama I genome as a reference. A maximum-likelihood phylogeny was inferred using IQ-Tree 2 (v2.0.4, 43). IQ-Tree’s built-in ModelFinder (44) module was used to select the best model for each dataset. Up to 2,000 bootstrapping steps were conducted, using the built-in Ultrafast bootstrap module (45). The best maximum likelihood tree for each dataset was output in Newick format, which was then visualised using Microreact (46) or iTOL (47).

## 7. Results

### Closed genome sequences were assembled for all isolates

In total, 6 MinION R9.4 RevD flow cells were used to sequence the 66 New Zealand isolates. The amount of usable data per flow cell, after demultiplexing with Deepbinner and Guppy, varied from 3.94 to 12.76 Gb. This corresponded to coverage between 42X and 484X per isolate, with a mean coverage of 207X. The overall mean read length across all sequencing runs was 6,282 bp. The individual run mean read lengths ranged from 4,908 to 7,917 bp.

The coverage and read lengths were sufficient to assemble closed genome sequences for all isolates, 63 in hybrid with Illumina data, and 3 using nanopore data alone (full details shown in **Supplementary Table S1**).

### 19 different genome arrangements were observed in the 66 New Zealand isolates

Closed genome sequences were produced for all 66 New Zealand isolates, and the genome arrangement of these closed sequences was investigated using progressiveMauve. The isolates were grouped based on shared arrangement, resulting in 19 different arrangement groups. Ten isolates displayed “singleton” arrangements which were not shared with any other isolate (**Supplementary Figure S1**); the remaining 56 isolates shared nine arrangements between them (shown in **Figure 1a**). As seen in Weigand, Peng (16), the rearrangements generally displayed a pattern of symmetrical inversions around the origin of replication.

**Fig. 1.**
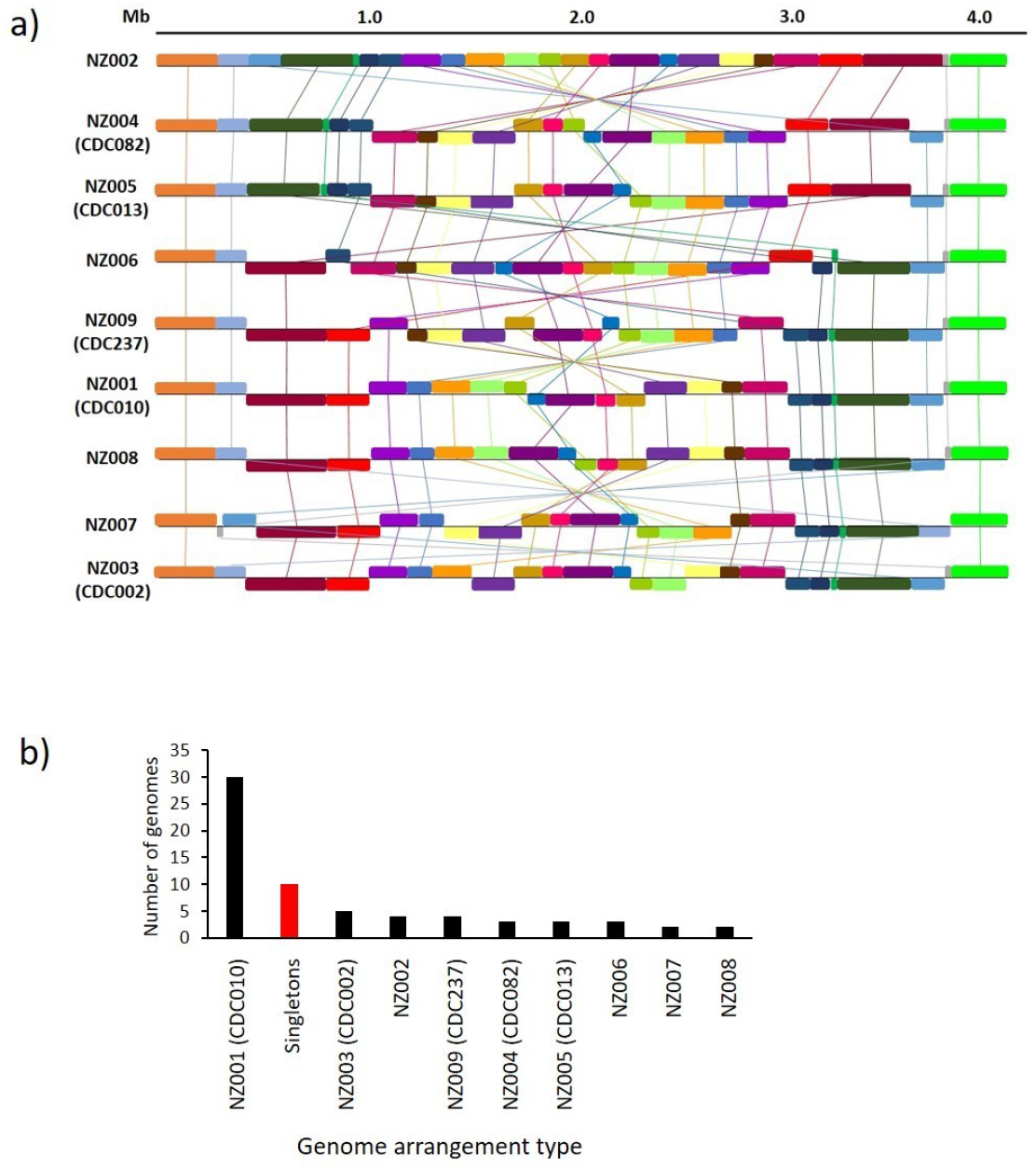
Genome structures of New Zealand isolates. a) 56 of the 66 isolates could be grouped into nine shared genome arrangements. The differences between the arrangement types tended to be inversions (often large) around the central region of the chromosome. Five of the nine arrangement types were found to be congruent with CDC structures defined in Weigand et al. 2017 and/or Weigand et al. 2019 (16, 17). One of the New Zealand isolates with a “singleton” arrangement (NZ48) was also found to be congruent with CDC046 (not shown), whilst another was congruent with CDC Cluster-BP-23 (NZ10). b) 30 New Zealand isolates shared the same arrangement type (NZ001/CDC010). Ten isolates had “singleton” arrangements shared with no other isolate (although one of these arrangements was found to be congruent with CDC046). The remaining 26 isolates were split more evenly between the other eight arrangement types.

One representative of each arrangement group, including each singleton, was aligned using progressiveMauve against representatives of each of the arrangement types defined in Weigand, Peng (17) and Weigand, Peng (16) (the representatives are listed in **Supplementary Table S3**). This revealed that six of the New Zealand arrangements were congruent with Weigand arrangement types, including two of the New Zealand singletons, NZ10 and NZ48, which were congruent with the CDC Cluster-BP-23 and CDC046 arrangement types, respectively. 30 New Zealand isolates shared the same arrangement type, CDC010. The remaining isolates were spread more evenly in smaller groups across the other eight arrangement types, as shown in **Figure 1b**. Full details of the grouped arrangement types are given in **Supplementary Table S5**. Arrangement type CDC010 was the fifth most commonly seen arrangement type in Weigand, Peng (17), after CDC237, CDC002, singletons and CDC013; CDC237, CDC002 and CDC013 were also observed here.

### Shifts in allelic profiles of New Zealand isolates have paralleled those seen elsewhere

MLST was used to identify which alleles of *ptxP, ptxA, prn, fim2* and *fim3* were present in the 63 New Zealand isolates for which Illumina data was available, using allele definitions from PubMLST or as defined in Bart, Harris (7). The results are shown in **Figure 2**. The most common allelic profile in New Zealand in the WCV era, particularly before the 1990s, was *ptxP1-ptxA1-prn1-fim2-1-fim3-1* (75% of 1980s strains, n=8). This then shifted gradually to *ptxP3-ptxA1-prn2-fim2-1-fim3-1* (90.3% of 2010s strains, n=31) throughout the 1990s, 2000s and 2010s. A brief increase in the circulating proportion of *fim3-2* was seen, although this decreased again by the 2010s.

**Fig. 2.**
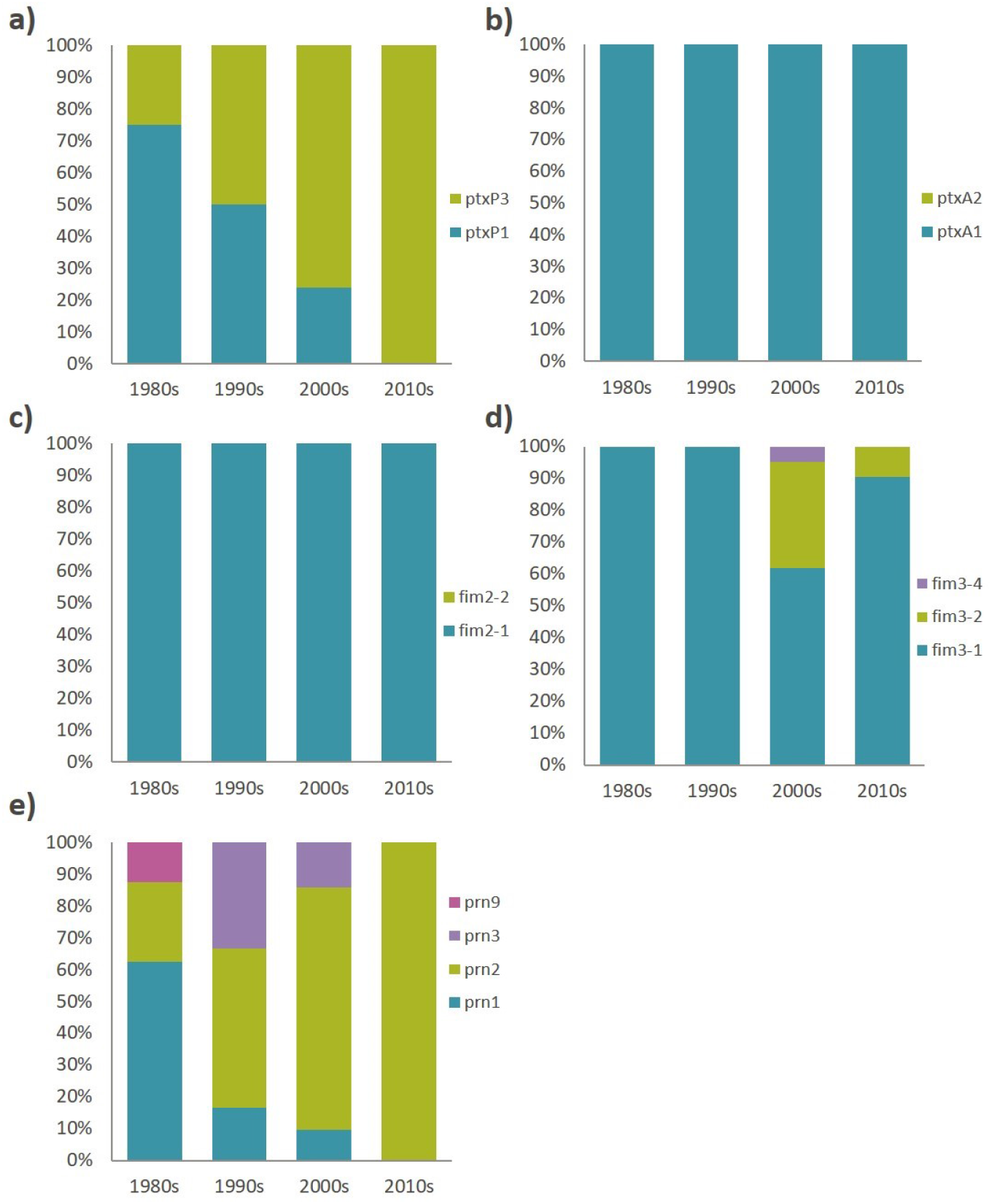
The changing allelelic profile of New Zealand strains from 1982 to 2018. Some of the genes involved in the ACV have undergone noticeable shifts globally since the switch from WCV to ACV in the late 1990s and early 2000s, in addition to a rapid shift from *ptxP1* to *ptxP3* in the 1980s and early 1990s. a) Shows that the shift from *ptxP1* to *ptxP3* also occurred in New Zealand, but *ptxP1* alleles continued to circulate in New Zealand until the 2000s. b) and c) show that, in New Zealand, one allele each is circulating for *ptxA* and *fim2*. d) Shows a brief increase in the frequencies of non-*fim3-1* alleles. e) Shows a rapid increase in the prevalence of *prn2* since the 1990s; no other *prn* alleles appear to be present in the population currently.

During the 1990s and 2000s, a number of strains carrying the *prn3* allele circulated, partly related to the 2004-2006 outbreak. However, by the 2010s, only *prn2* was circulating. Additionally, 44.4% (n=9) of strains circulating during the 2004-2006 outbreak carried the *ptxP1* allele, which had largely been replaced by *ptxP3* throughout the 1990s. Interestingly, three of the four *ptxP1* strains isolated during the 2004-2006 outbreak also carried *prn3*, and shared the same genomic arrangement (arrangement type NZ002). Two of these strains were isolated within a month of each other in the same region (NZ35 and NZ36, Midcentral), so may have been directly related. However, the third strain (NZ37) was isolated nearly a year later, in a different region (Capital and Coast). This suggests that *ptxP1-prn3* may have been a common allelic profile during that outbreak. No other similar phenomena were observed in the 2008-2011, 2012 or 2017-2018 outbreaks.

### 35% of all strains, and 89% of strains isolated since 2012, were predicted to be pertactin deficient

The closed genome sequences were annotated using Prokka, and the resulting annotations were screened for the presence/absence of *prn*, *fhaB* and *ptxA-E*, as well as for the insertion of IS *481*, IS *1002* or IS *1663* into the same genes.

34.8% (23/66) of all strains were found to have a copy of IS *481* within their *prn* gene, which is known to cause pertactin deficiency. No pre-2012 strain (n=39) was found to have this insertion, whilst 85.2% (23/27) of all strains isolated between 2012 and 2018 had it. There are three potential IS *481* insertion sites within the *prn* gene, at 240 bp, 1,613 bp, and 2,735 bp (41). An IS *481* insertion occurred at the 1,613 bp site in 22/23 of the New Zealand strains with an IS *481* insertion in *prn*. The remaining strain had IS *481* inserted at position 240. Full details of the *prn* mutation in each strain are shown in **Supplementary Table S1**.

One additional strain (NZ63) from 2017 was found to have a mutation previously identified in Weigand et al. (2017) as causing deficiency, a change from C to T at position 223, resulting in a premature stop codon. In total, therefore, 88.9% (24/27) of strains isolated in 2012 or later are predicted to be pertactin-deficient. This suggests that pertactin deficiency was uncommon in New Zealand *B. pertussis* strains until 2012, and has rapidly become prevalent since then.

No FHA or PT deficiency caused by IS insertion was predicted in any strain.

### New Zealand isolates mainly cluster according to allelic profile rather than outbreak or year of isolation, with some notable exceptions

A phylogeny was inferred using 325 core SNPs for 63 New Zealand isolates (**Figure 3**). The *ptxP1* strains clustered separately from the *ptxP3* strains. All *non-prn2* strains were contained in the *ptxP1* cluster. All predicted PRN-deficient strains occurred in the *ptxP3* cluster, with all but one of them clustered on the same sub-branch within the *ptxP3* cluster. However, the different deficiency-causing *prn2* mutations did not cluster together within this sub-branch (for example, the IS *481*-1613fwd and IS *481*-1613rev strains did not cluster separately), suggesting each mechanism for PRN deficiency has arisen on multiple occasions.

**Fig. 3.**
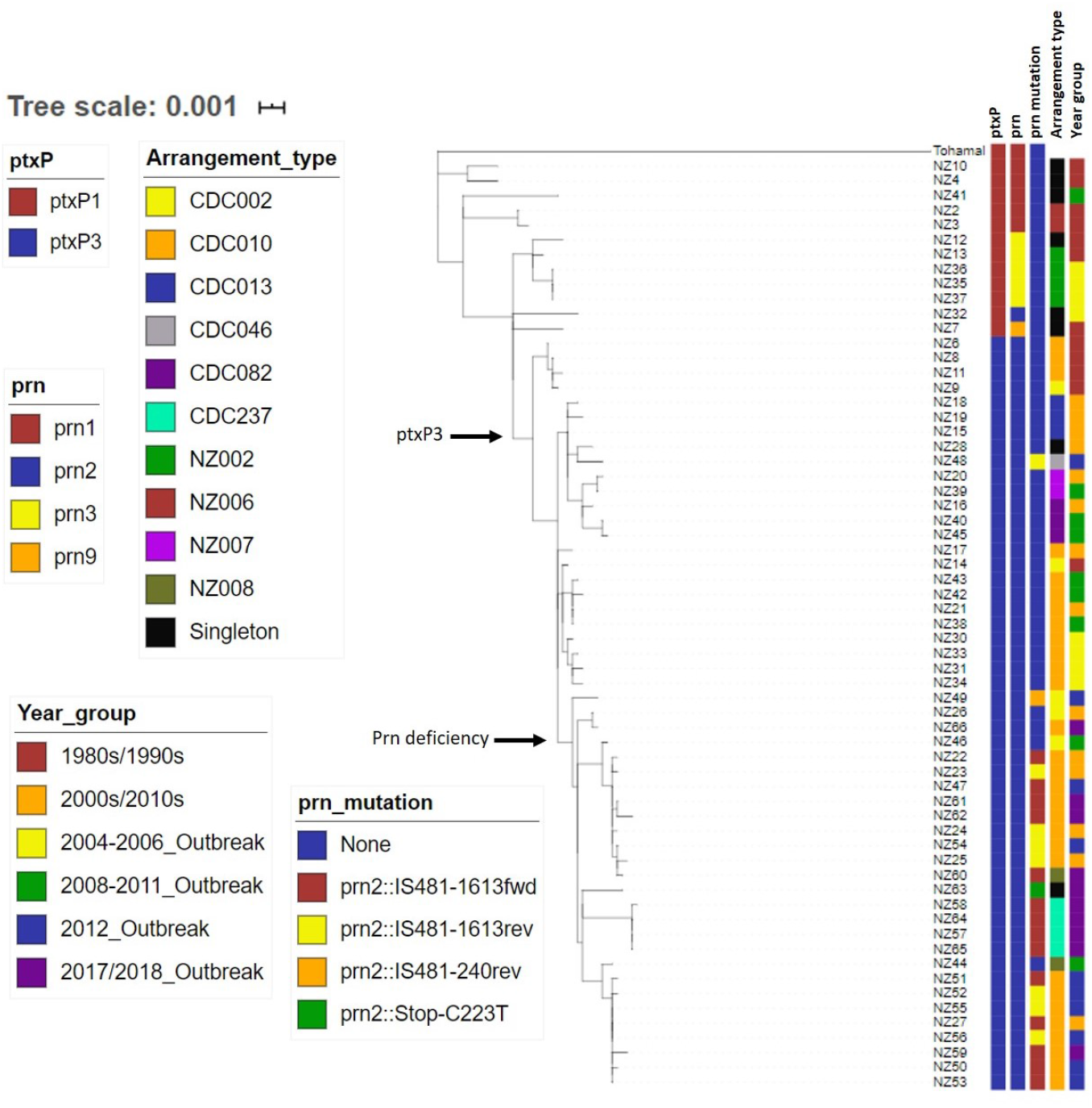
Phylogenetic tree showing the evolutionary relationships of the New Zealand strains sequenced here, in the context of allele type, Prn deficiency, genomic arrangement and period of isolation. Snippy was used to identify variants between the NZ strains and the reference, Tohama I, as well as defining a core genome. IQ-Tree2 was used to infer a maximum-likelihood phylogeny, which was then displayed using iTOL (interactive Microreact tree also available at https://microreact.org/project/JG7I7MBBV). Key branches are highlighted. All ptxP3 strains are contained within one major branch, whilst a slightly smaller sub-branch contains all but one of the predicted Prn-deficient strains.

Most arrangement types were spread relatively evenly throughout the phylogeny; however, strains with the NZ002 arrangement type clustered, including the three highly similar strains from the 2004-2006 outbreak. These strains (NZ35, NZ36 and NZ37) were identical in terms of core genome SNPs; comparing the NZ36 and NZ37 Illumina reads to the hybrid assembled NZ35 genome showed that NZ35 and NZ36 were also identical in their whole genomes, whilst NZ37 was only one SNP different.

Likewise, strains with the arrangement type CDC237 clustered, and all were isolated during the most recent (2017-2018) outbreak, representing 40% (n=10) of all strains sequenced from that outbreak. These four strains (NZ57, NZ58, NZ64 and NZ65) also shared the same mutation in their *prn2* gene (IS *481*-1613fwd), and three of them (NZ57, NZ64 and NZ65) were identical in terms of core genome SNPs. Comparing the Illumina reads for the four strains to the hybrid assembled whole genome sequence of NZ64 showed that the cluster is separated by only 1-2 SNPs across the whole genome, with NZ57 and NZ65 being identical across the whole genome. Two of the strains (NZ57 and NZ58) were isolated in the Southern region within the same two-week period, whilst the other two strains (NZ64 and NZ65) were isolated in the South Canterbury region several months later, but again within two weeks of each other. 70% (n=10) of the strains isolated during the 2017-2018 outbreak period clustered, regardless of genomic arrangement, suggesting that the many of the strains circulating during that outbreak were closely related.

Other than the 2004-2006 and 2017-2018 outbreaks, strains did not tend to cluster according to the outbreak or year in which they were isolated. On the whole, many New Zealand strains were very closely related, including a large group of eight strains (NZ51, NZ52, NZ55, NZ27, NZ56, NZ59, NZ50 and NZ53) collected throughout the 2000s/2010s, which were either identical or separated by only one or two SNPs in their core genome. Variant calling with Snippy, with the hybrid assembled NZ56 genome as a reference and the Illumina reads for the other seven strains, showed that six of the strains (NZ56, NZ51, NZ52, NZ53, NZ55 and NZ27) were separated by no more than three SNPs across their whole genome, whilst the remaining two strains (NZ59 and NZ50) were eleven and six SNPs different from NZ56, respectively.

### New Zealand isolates cluster phylogenetically with global strains according to allelic profile, with some clusters of closely related isolates

To determine whether the New Zealand strains were genetically distinct from those circulating elsewhere in the world during 1982-2018, a further phylogeny was inferred from the 63 New Zealand strains and 73 global strains from the same period, using 511 core SNPs (**Figure 4Fig.)**. Again, all *ptxP1* strains clustered on the same branch, and all *ptxP3* strains clustered on a separate, larger branch. In general, strains isolated earlier (for example, in the 1980s) clustered at one end of the tree, whilst strains isolated later (for example, the 2010s) clustered at the other end.

**Fig. 4.**
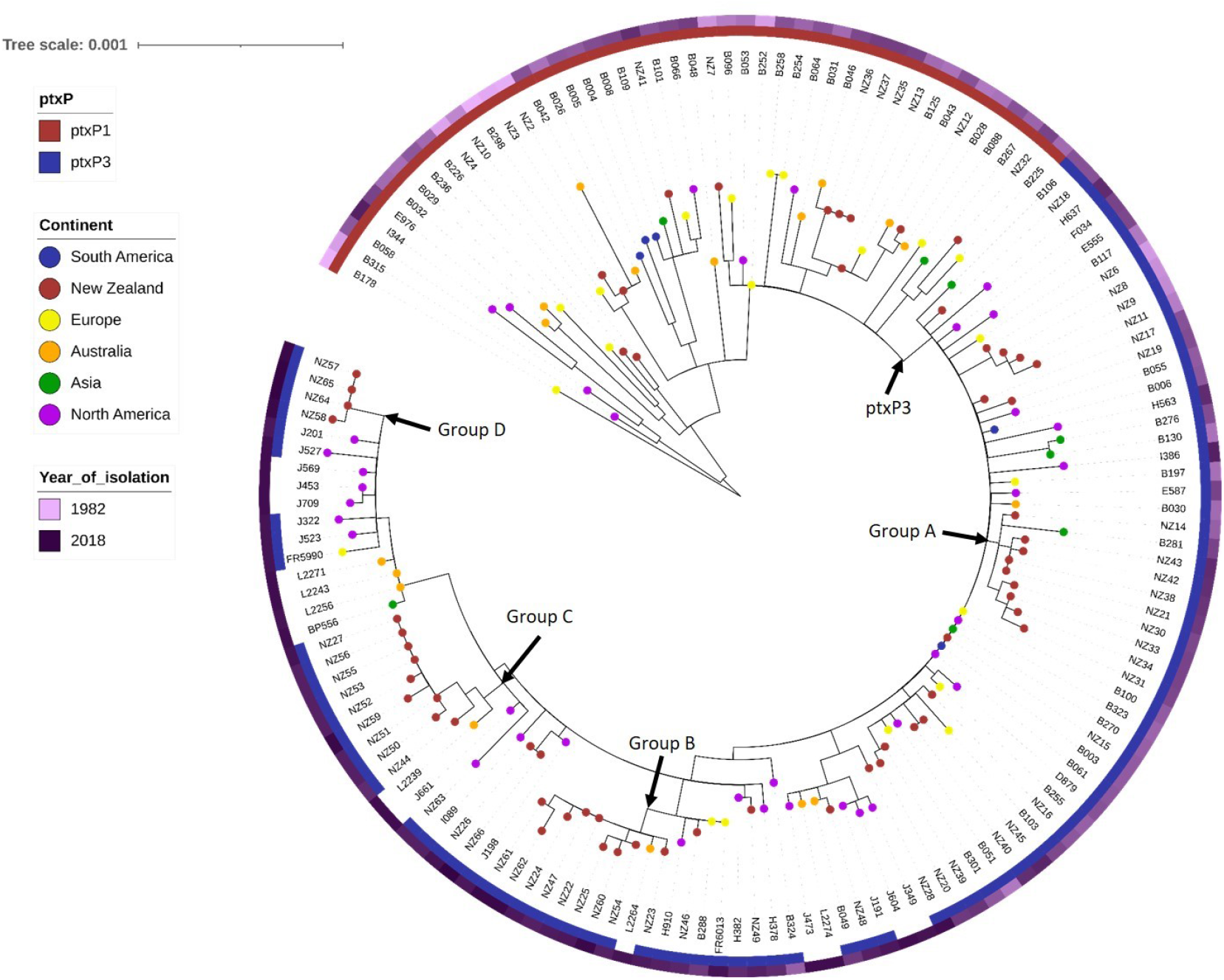
Phylogenetic tree showing the evolutionary relationships of the New Zealand strains compared with a selection of global strains from the same time period. Variants between the selected strains and the Tohama I reference were called, and a core genome was defined, using Snippy. A maximum-likelihood phylogeny was inferred using IQ-Tree2, and iTOL was used to display the resulting tree. Like the New Zealand strains alone, all strains carrying the *ptxP3* allele* are found on the same major branch, separate from the *ptxP1* strains. Several clusters containing highly similar strains, almost all from New Zealand, are indicated. *Allele information was not available for some of the more recent global strains.

During periods for which sequencing data was available from many countries, little geographical clustering was seen. For recent years (since 2014), most of the available sequencing data was for strains isolated in the USA and sequenced by the CDC; this gives the impression of geographical clustering, but is more likely simply due to under-sampling of strains from other countries.

However, certain New Zealand strains do group separately from the majority of the global strains. Groups A-C each contain ten isolates (nine which were isolated in New Zealand, and one which was isolated in either Australia (Groups B and C) or Asia (Group A)), which are clustered on their own sub-branches of the global tree, separate from the other groups and from the rest of their closest global neighbours. The isolates contained in each group are all strikingly similar to each other in terms of allelic profile, genomic arrangement, presence/absence of specific *prn2* deficiency-causing mutations and, where data was available, serotype. Perhaps surprisingly, these samples were not all isolated in the same location, and nor were they always isolated around the same time (although all were isolated since 1999); in Group A, for example, the oldest isolate was from 1999, whilst the newest was from 2010. Further details of Groups A-C are shown in **Supplementary Table S6.**

Group D is a smaller cluster of four strains (NZ57, NZ58, NZ64 and NZ65) which was previously identified in the New Zealand phylogeny. As mentioned previously, these four strains are separated by at most one SNP across their whole genomes, and two (NZ56 and NZ53) are identical. All have the RD4 deletion and the IS *481*-1613fwd *prn2* mutation. They all also have the CDC237 arrangement type, and were isolated in two contemporaneous and co-located pairs.

### Isolates circulating during outbreak years appear to be more closely related to each other than outbreak strains in other countries

Groups A-D in the global phylogeny suggest that certain strains circulating in New Zealand in the 2000s and 2010s were more closely related to each other than those circulating in other countries, particularly during New Zealand whooping cough outbreak periods. To investigate whether this effect was simply due to oversampling of New Zealand strains in the global phylogeny, smaller phylogenies were constructed for each of the four post-2000 New Zealand outbreaks, with additional global strains included for each outbreak period.

**Figure 5** shows the focussed phylogenies for a) the 2004-2006 outbreak and b) the 2008-2011 outbreak. The New Zealand strains are spread across several different branches in the 2004-2006 phylogeny, which was constructed from 378 core SNPs. Nonetheless certain strains are closely related: NZ35, NZ36 and NZ37, as mentioned previously, are identical in terms of core SNPs, and were not all isolated at the same time or location. However, on the same branch as these three New Zealand strains are two Australian strains. These Australian strains are identical to each other and differ from the New Zealand strains by only two core SNPs. Similarly, NZ32 and L508 (also from Australia) are identical to each other in their core genomes, as are two other Australian strains, B048 and L518. Overall, this phylogeny suggests that certain strains circulating during the 2004-2006 outbreak were clonal in both New Zealand and Australia, but the outbreak was not caused by a single strain in either country.

**Fig. 5.**
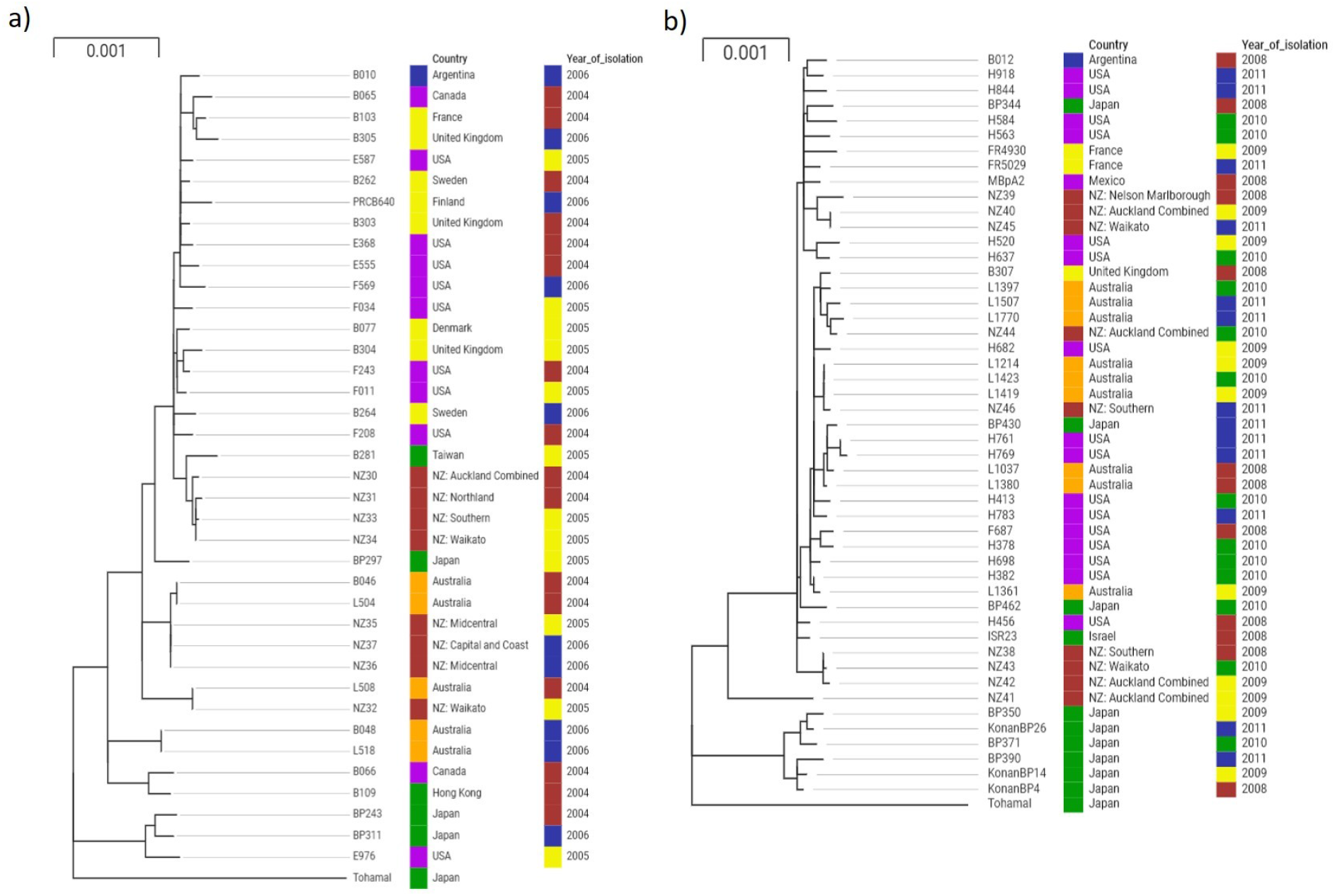
Phylogenetic trees showing the evolutionary relationships between strains isolated in New Zealand during the a) 2004-2006 whooping cough outbreak and b) 2008-2011 whooping cough outbreak, compared with global strains isolated during the same time periods. Variants were called and core genomes defined using Snippy, then phylogenies were inferred using IQ-Tree2 and visualised using Microreact (interactive trees available at https://microreact.org/project/8F3Ilz2Bi and https://microreact.org/project/UWL4KNRaN)

The picture is less clear for the 2008-2011 phylogeny, which was inferred from 408 core SNPs. Again, some of the New Zealand and Australian strains cluster separately from strains from elsewhere. Three Australian strains (L1214, L1423 and L1419) have identical core genomes and one New Zealand strain (NZ46) differs by only one SNP. Likewise, there are three New Zealand strains from different regions (NZ38, NZ43 and NZ42) which are, at most, only one SNP apart. However, on the whole, the New Zealand strains from the 2008-2011 outbreak are less closely related to each other than those from the 2004-2006 outbreak, and there is less of a clear similarity with the Australian strains.

**Figure 6** shows the focussed phylogenies for the a) 2012 outbreak and b) the 2017-2018 outbreak. The 2012 phylogeny, inferred from 320 core SNPs, shows that 60% (n=10) of the New Zealand strains (NZ52, NZ50, NZ53, NZ51, NZ55 and NZ56) form a clear group, again with some Australian strains (L1661, L1779 and L1780). Four of the New Zealand strains have identical core genomes, and the others differ by one SNP each. The Australian strains are not identical to each other, but each is only two SNPs different from the New Zealand strains. The remaining New Zealand strains are spread throughout the rest of the phylogeny, with strains isolated in the UK or USA.

**Fig. 6.**
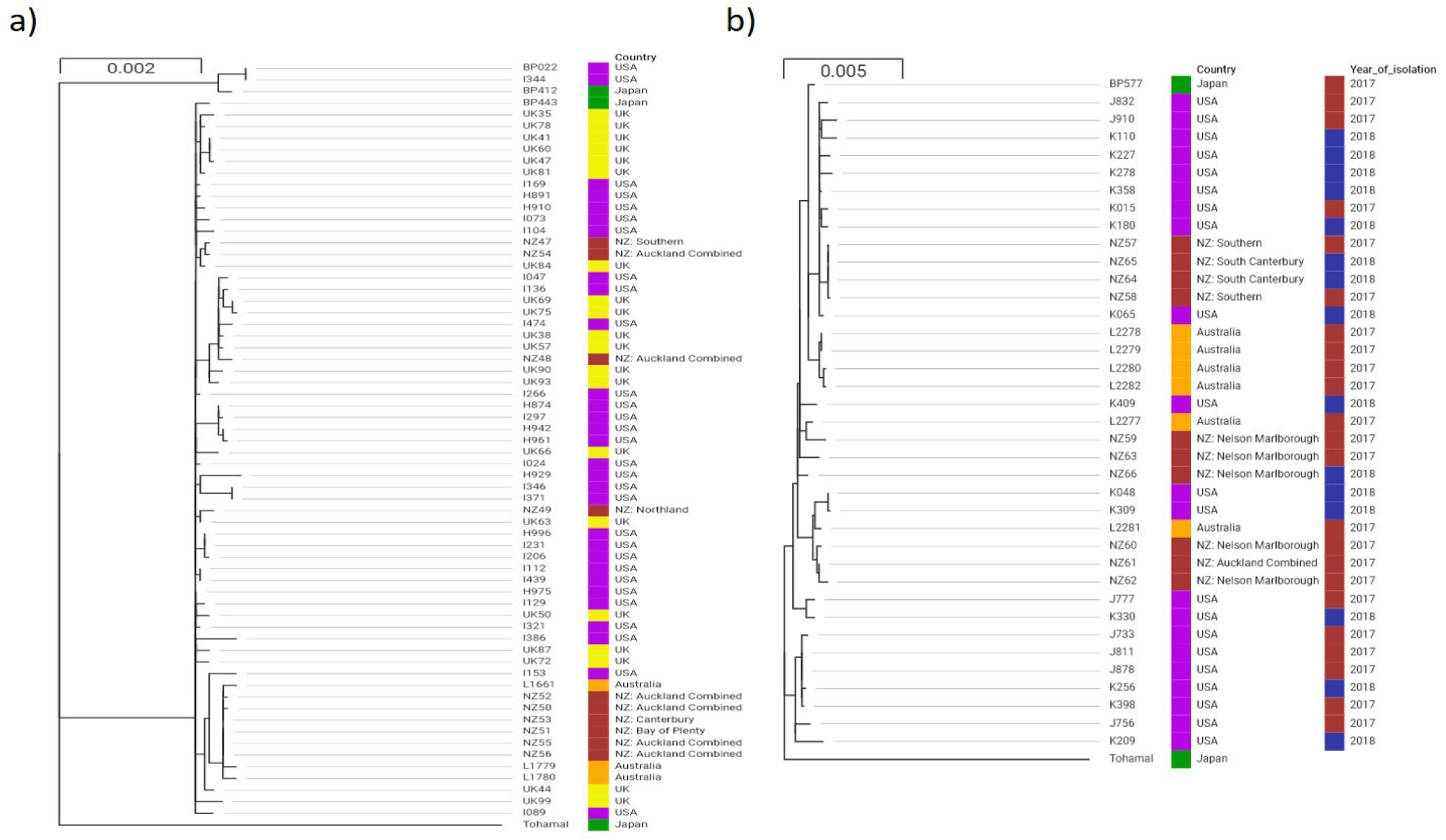
Phylogenetic trees showing the evolutionary relationships between strains isolated in New Zealand during the a) 2012 whooping cough outbreak and b) 2017-2018 whooping cough outbreak, compared with global strains isolated during the same time periods. Variants were called and core genomes defined using Snippy, then phylogenies were inferred using IQ-Tree2 and visualised using Microreact (interactive trees available at https://microreact.org/project/yR6WukJhF and https://microreact.org/project/tbl0zUnck)

Finally, the phylogeny showing strains from the most recent outbreak, which was constructed from 323 core genome SNPs, indicates a slightly different picture again. As previously mentioned, four of the New Zealand strains (NZ57, NZ65, NZ64 and NZ58) have identical core genomes; these strains cluster on their own branch. The remaining New Zealand strains, however, are less similar to each other, and cluster on branches with strains from Australia and/or the USA. At the same time, several USA strains (J733, J811, J878, K256 and K298) isolated over the same period are identical or only a single SNP different in their core genomes.

## 8. Discussion

New Zealand strains were investigated here for a number of reasons. Historically, vaccine uptake in New Zealand was lower than in many other countries although efforts to monitor and increase rates of uptake since 2008 mean that vaccine coverage is now comparable to that in the UK. Despite the increasing vaccine coverage, whooping cough incidence has remained higher in New Zealand than in most other countries, including the USA and UK. An outbreak occurred in New Zealand from 2017 to 2019, for example - a period when no other countries noted a similar increase in cases. Although isolates have been collected and stored by New Zealand’s ESR since the 1980s, no sequencing of any *B. pertussis* from New Zealand had previously been conducted. Therefore, the genomes of 66 isolates from between 1982 and 2018 were sequenced here, using hybrid nanopore and Illumina sequencing, to determine the following: whether global genomic trends observed in *B. pertussis* had been delayed by New Zealand’s slower vaccine uptake; whether the strains circulating during whooping cough outbreaks were polyclonal, as seen in recent outbreaks in the UK and USA; and whether the consistently higher incidence of whooping cough in New Zealand could potentially be explained by the circulation of a unique hypervirulent strain.

### Allelic profile and antigen deficiency trends in New Zealand generally match those observed elsewhere in the world

Analysis of the allelic profile of the ACV-related genes in the New Zealand isolates reveals a pattern in the New Zealand strains similar to that seen in Bart et al.’s landmark 2014 study of global strains throughout the 20^th^ and early 21^st^ century, as well as other studies of how *B. pertussis* populations have changed over recent years (2, 7, 48, 49). The most common global allelic profile during the “WCV era” (defined as 1960-1995) was *ptxP1-ptxA1-prn1-fim2-1-fim3-1* (7). The most common allelic profile of the New Zealand strains from 1982 to 1995 was the same, and was present in 50% (n=12) of the strains screened; the other 50% of strains were split between those carrying *ptxP3*, and different *prn* alleles. In the Bart et al. “WCV/ACV era” and “ACV era” (post-1995), the most prevalent allelic profile shifted to *ptxP3-ptxA1-prn2-fim2-1-fim3-1*. An increased frequency of strains carrying the newer *fim3-2* allele was also noted (from <1% frequency in the WCV era to 37% in the ACV era), with *ptxP3-ptxA1-prn2-fim2-1-fim3-2* being observed as the dominant profile in the late 2010s (50). Again, the most common allelic profile seen in the New Zealand strains throughout the period matches that seen elsewhere in the world.

Noted shifts, such as *ptxP1* to *ptxP3* and *prn1* to *prn2*, appear to have happened in New Zealand on similar timescales to the rest of the world. Interestingly, although the progression of *fim3-2* in the New Zealand strains reflects that seen in Bart, et al. 2014 (7) throughout the WCV and early ACV eras, rising from 0% of strains in the 1980s and 1990s to 30.4% (7/23 strains) in the 2000s, a decline in the frequency of this allele is apparent in the strains from the 2010s (to 9.7%, 3/31 strains, **Figure 2Fig.)**. The most recent strains in the Bart et al. screen were isolated in 2010; it is therefore possible that a similar decrease in prevalence would be seen in more recent global strains, rather than being a New Zealand-specific phenomenon. This is supported by studies such as Bowden, Williams (6), which note a re-emergence of the *fim3-1* allele since 2010.

Another shift observed globally in the ACV era has been the rapid increase in strains which do not express functional pertactin protein. In Australia, for example, the percentage of PRN-deficient strains increased from 5% to 78% over the period 2008 to 2012 (3). A study of strains from European countries isolated between 1998 and 2015 showed a clear correlation between how early each country introduced the ACV and the current proportion of PRN-deficient strains (12). The ACV was introduced in New Zealand in 2000, several years before some European countries, including the UK. Although the first PRN-deficient strains did not appear in the New Zealand cohort studied here until 2012, later than in some countries, the proportion of PRN-deficient strains rapidly increased; 88.9% of strains since 2012 are PRN-deficient, including 90% of those isolated in 2017 and 2018. This frequency is higher than for any of the European strains included in Barkoff et al.’s study (12), although it is similar to the frequencies observed in Australia (78%) and the USA (85%) (3, 11). The majority of the predicted PRN-deficient New Zealand strains (95.8%, n=24) have an insertion of IS *481* in their *prn* gene, and one (NZ63) possesses a SNP which results in a premature stop codon. Other mechanisms for PRN deficiency have been observed in other studies (for example, 41), such as a commonly observed inversion of 22 kb containing the *prn* promoter, but were not observed here. Yet further mechanisms for PRN deficiency may not have been identified from sequence alone as, in previous studies, the mechanism for deficiency has not always been identifiable. The percentage of PRN-deficient strains in New Zealand may therefore be even higher than predicted. Expression or non-expression of antigens such as PRN, PT and FHA is often tested *in vitro*, for example by western blot, but only heat-killed cells were available here. Nonetheless, it should be noted that the use of hybrid genome assembly allowed the prediction from sequence alone of PRN deficiency caused by the most common deficiency mutations.

Overall, New Zealand’s historically lower coverage of the whooping cough vaccine does not seem to have significantly delayed the most commonly observed recent genomic changes, unlike countries which have been slower to take up the vaccine, such as China (48).

### The *B. pertussis* strains circulating in New Zealand appear to be more clonal than in many other countries

A variety of global *B. pertussis* phylogenies have been produced over the last decade, including the landmark Bart et al. 2014 study (7). Some of the key patterns from these phylogenies, for example the clear branching of *ptxP1* from the *ptxP3* strains, were also observed in the global phylogeny here (**Figure 4**), with the New Zealand isolates clustering with the sequences from the rest of the world. Another discovery of Bart et al. was that there was very little geographic clustering of strains; strains from all around the world were spread throughout the phylogeny. Likewise, Sealey’s (51) investigation of UK strains showed little geographic specificity. In addition, studies of specific outbreaks in the UK and USA showed that strains circulating during outbreaks were polyclonal, and not characterised by a single outbreak strain (2, 6). Conversely, an outbreak in Australia in the late 2000s and early 2010s was found to be caused primarily by the circulation of a group of highly related strains (4).

Although some of the New Zealand isolates cluster throughout the global phylogeny along with strains from other countries, some notable groups (labelled Groups A-D in **Figure 4**) of highly similar New Zealand isolates stand apart. Groups A-C each contain nine New Zealand strains and one strain each from either Australia (Groups B and C) or Asia (Group A), all of which are identical in terms of core genome, or only one or two core genome SNPs different. Interestingly, the highly similar strains were not always isolated during the same year, often not even during the same outbreak, and all are from the post-2000 era. In each group, the strains are not only similar in terms of core genome SNPs, but also in serotype (where tested), PRN deficiency and, usually, genomic arrangement.

Group D contains four identical New Zealand strains, all from the 2017-2018 outbreak. The four strains can be separated into two pairs which were each isolated in the same region and within the same timeframe. Although this suggests that the similarities observed in this cluster could be explained by transmission of the same strain between patients, the fact that the two pairs were isolated in different regions, and several months apart, could also indicate that the 2017-2019 outbreak may have in part been the result of the circulation of a particularly virulent strain around the country. This theory is challenged by the fact that the other six isolates sequenced from the outbreak did not cluster with Group D. Sequencing of more strains from the currently under-sampled outbreak could allow confirmation or rejection of the theory.

In case the clustering of New Zealand strains on the global tree was due to overrepresentation of New Zealand sequences compared to any other country, further phylogenies were constructed for the four major outbreaks since 2000, each including a greater number of global strains from the outbreak period. As New Zealand’s closest neighbour, it seemed possible that strains from Australia would show most similarity to New Zealand strains, so extra Australian strains were included where possible. In each of the trees, strains from Australia did indeed cluster most closely with the New Zealand strains, particularly during the 2004-2006 and 2012 outbreaks. Additionally, as seen in the global tree (**Figure 4**), some of the Australian strains had highly similar or identical genetic characteristics to some of the New Zealand strains. Three closed *B. pertussis* genome sequences were also available for Australian isolates, sequenced by Fong et al. (52); the genomic arrangements of these three strains were compared to each of the identified New Zealand arrangement types, including the singletons, to further investigate the apparent close similarities between isolates from the two countries. However, only one of the Australian isolates shared an arrangement with any of the New Zealand isolates: CIDM-BP2 and NZ48 both belonged to cluster CDC046, first identified by Weigand et al. (16, 17). Like eight of the New Zealand isolates, two of the Australian isolates had singleton arrangements, not yet seen in any other isolates globally. Seemingly, whilst some isolates from Australia are highly similar to those from New Zealand, both countries also have circulating isolates which are more geographically unique.

Whilst not indicating that the outbreaks in New Zealand have been caused by a single hypervirulent strain, the phylogenies do suggest that there may be a higher than usual proportion of very similar or identical strains circulating in New Zealand. These strains have been isolated during non-outbreak periods, although more often during outbreaks. It is possible that these clusters of strains are more virulent than most *B. pertussis*, as suggested by their overrepresentation during outbreak periods. Their circulation during non-outbreak years could therefore help to explain New Zealand’s seemingly permanent higher incidence of whooping cough than elsewhere in the world. However, such a phenomenon has not been observed in *B. pertussis* before, and there is no genetic evidence here for enhanced virulence, with the clusters all showing allelic profiles and other characteristics which are in line with the global trends.

The global and outbreak phylogenies are based on shared core genome SNPs compared to the *B. pertussis* reference strain, Tohama I. Thus, they do not include all of the SNPs in each strain, and seemingly identical strains may therefore not have been identical in their whole genomes. This was tested by comparing the Illumina reads from clustered strains to a single hybrid assembled whole genome belonging to that same cluster. This revealed that, in some cases, the strains which appeared to be identical in their core genomes were indeed identical across their whole genomes, whilst others differed across their whole genomes, but only by up to 11 SNPs (as in the case of NZ56 and NZ59). Even those strains which differed slightly across their whole genomes shared other similarities identified here, such as genome arrangement, serotype and *prn* mutation. The global and outbreak trees therefore show evidence of some geographic specificity, and that many of the New Zealand strains are more closely related to each other than to strains from other countries except, perhaps, Australia. Not all New Zealand strains fall into clustered groups, however; some occur throughout the phylogeny in clusters with strains from other countries, as seen with the UK strains in Sealey, 2015 (2).

The results presented here contribute to a growing body of knowledge on *B. pertussis* genomics, revealing changes that have occurred over time, influenced by the introduction and alteration of vaccines, as well as representing the first New Zealand *B. pertussis* isolates to be sequenced. This study also shows for the first time the utility of nanopore sequencing in conducting rapid and affordable *B. pertussis* strain screens, in combination with Illumina sequencing. The benefits of closed genome sequences for comparative genomics are demonstrated by the ability to group isolates not only by allelic profile, but also by genomic arrangement type. With a growing number of closed genome sequences to which the arrangement of future isolates can be compared, along with evidence that differences in genomic arrangement may influence the phenotypic behaviour of isolates (15), the ability to easily produce closed genome sequences is becoming increasingly useful. Finally, we were able to use the closed genome sequences to predict pertactin deficiency; in light of globally increasing antigen deficiency, this ability, particularly in the absence of live *B. pertussis* cells, could be especially beneficial in adding to our understanding of this ongoing trend.

## 9. Author statements

### 9.1 Authors and contributors

Conceptualization: All authors; Data curation: NR; Formal analysis: NR; Funding acquisition: AP & SB; Investigation: NR, HD, JM, MS & AT; Methodology: NR; Project administration: All authors; Resources: HD, JM, MS & AT; Supervision: AP & SB; Validation: NR; Visualization: NR; Writing - original draft: NR; Writing - review & editing: AP, AT & SB.

### 9.2 Conflicts of interest

N.R. was part-funded by Oxford Nanopore Technologies to conduct PhD research. No other conflicts of interest exist.

### 9.3 Funding information

This work was funded by the University of Bath and Oxford Nanopore Technologies. The New Zealand isolates were originally collected, stored, prepared and shipped by the Institute of Environmental Science and Research, funded by the New Zealand Ministry of Health with the cooperation of the local diagnostic laboratories.

## 9.4 Acknowledgements

The authors are grateful to the Institute of Environmental Science and Research and the New Zealand Ministry of Health for providing the isolates investigated here, and to Oxford Nanopore Technologies for part-funding NR’s PhD studentship. We also continue to be thankful for the fantastic bioinformatics resource, CLIMB (developed by the MRC, grant number MR/L015080/1), without which the data analysis undertaken here would not have been possible.

## Notes

https://www.ncbi.nlm.nih.gov/bioproject/PRJNA556877

### Summary of Updates

Fixed a typo in one of the authors' institutions.

