## Supplementary figures and images for "Comparative genomics of *Bordetella pertussis* isolates from New Zealand, a country with an uncommonly high incidence of whooping cough"

### Supplementary Figure S1

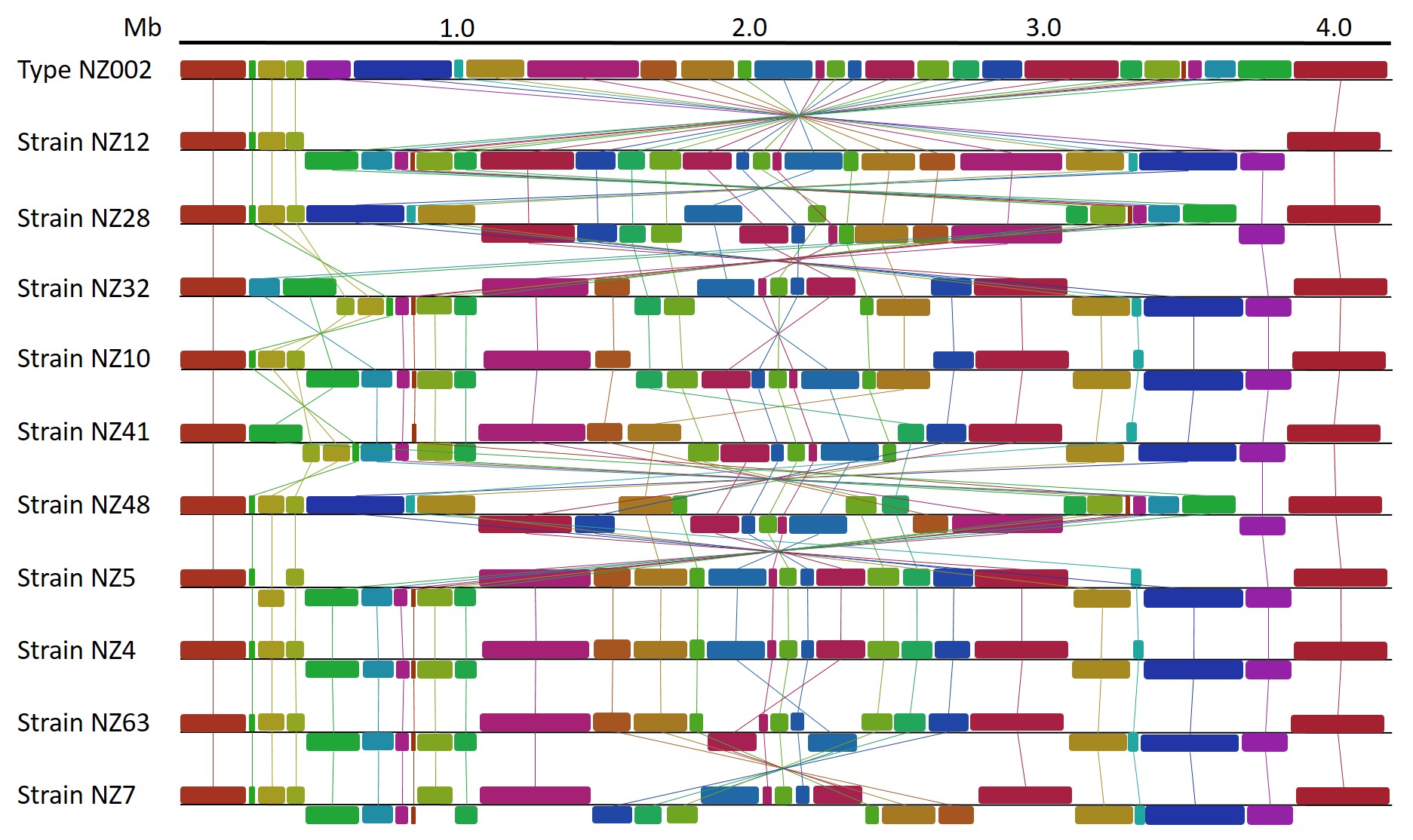
